# Redundant and distinct roles of two 14-3-3 proteins in *Fusarium sacchari*, pathogen of sugarcane Pokkah boeng disease

**DOI:** 10.1101/2023.04.10.536328

**Authors:** Yuejia Chen, Ziting Yao, Lixian Zhao, Mei Yu, Siying Qin, Chengwu Zou, Baoshan Chen

## Abstract

*Fusarium sacchari* is one of the most important sugarcane pathogens that causes Pokkah boeng disease (PBD) in China. 14-3-3 proteins have been shown to play vital roles in developmental processes in dimorphic transition, signal transduction and carbon metabolism in some phytopathogenic fungi, but were poorly understood in *F. sacchari*. In this study, two 14-3-3 protein-encoding genes, *FsBmh1* and *FsBmh2* in *F. sacchari*, were investigated. Although both *FsBmh1* and *FsBmh2* were expressed at vegetative growth stage, *FsBmh1* was repressed at sporulation stage in vitro. In order to clarify the roles of *FsBmh1* and *FsBmh2*, deletion mutants ΔFsBmh1 and ΔFsBmh2 were constructed. Phenotypic defects, including hyphal branching, hyphal septation, conidiation, spore germination and colony growth, were more severe in ΔFsBmh2 than in ΔFsBmh1, although virulence attenuation was observed in both mutants. To further explore the relationship between *FsBmh1* and *FsBmh2*, the combination of deletion and silencing (ΔFsBmh/sFsBmh) and overexpression (O-FsBmh) mutants were constructed and characterized. Compared to the single allele deletion, combinations of ΔFsBmh1/sFsBmh2 or ΔFsBmh2/sFsBmh1 showed more severe manifestations in general, suggesting a redundancy in function of the two 14-3-3 genes. Comparative transcriptome analysis revealed that more functional genes were affected in ΔFsBmh2 than in ΔFsBmh1. Redundancy in function between *FsBmh1* and *FsBmh2* suggests that 14-3-3 is vitally important for the organism and distinction in roles between the two isoforms may be resulted from the divergence in evolution. To the best of our knowledge, this was the first report on the distinct roles of 14-3-3 protein isoforms in a pathogenic fungus. Knowledge gained from this study should be of help to further understand the regulation mechanism of pathogenicity-related traits in fungal pathogens and for the development of new strategy for control of PBD in particular.

## Introduction

Sugarcane Pokkah boeng disease (PBD) is an airborne fungal disease caused by the *Fusarium* species complex (FSC) [1, 2]. The spores of the fungi are spread mainly through the air and rain to infect the sugarcane shoot tips, causing yellowing, crinkling, and twisting of the infected leaves and rotting of the tips [3]. In China, *F. sacchari* is the dominant pathogen causing PBD [4]. Though the disease has been investigated in varied aspects, e.g. identification of pathogens, screening of resistant germplasm resources [5], exploring the influence of environmental factors on the occurrence of PBD [6] and effectors [7], the pathogenic mechanism of *F. sacchari* is still unclear.

14-3-3 proteins are a protein family with low molecular weight of about 30 KDa, commonly found in eukaryotes [8, 9]. These proteins function by binding to and altering the activity, localization, and form of presentation of their target proteins to modulate a wide variety of cellular processes [10–13]. In the cell, 14-3-3 proteins are found to be in the form of homodimer or heterodimer [14]. Isoforms are common in 14-3-3 proteins, e.g., there are 15 isoforms in plant *Arabidopsis* and seven in mammals [15], but only one or two in fungal species [16, 17]. Although highly conserved between isoforms, different isoforms may have different affinities to bind their preferred targets and execute different functions [18].

In *S. cerevisiae*, 14-3-3-encoding genes *Bmh1* and *Bmh2* regulates vegetative growth and knockout of both genes would lead to lethal phenotype [19]. In human pathogen *Candida albicans*, *Bmh1* is essential for growth, morphogenesis, and pathogenesis [20]. In plant pathogen *U. maydis*, *Pdc1* encoding a homolog of *Bmh1* of the yeast, is involved in regulation of mycelial growth and virulence [21]. However, the function of the 14-3-3 proteins in *F. sacchari* remain uninvestigated. In this study, the function of 14-3-3 protein-encoding genes *FsBmh1* and *FsBmh2* in *F. sacchari* were investigated by a combinatory approach, gene deletion, silencing and overexpression. It was found that while both genes were required for virulence, *FsBmh2* played a major role in regulation of mycelial growth and conidiation in *F. sacchari*.

## Materials and methods

### Fungal strains and growth conditions

The *F. sacchari* wide-type strain CNO-1 and mutants including deletion mutants ΔFsBmh1 and ΔFsBmh2, a combination of deletion and silenced mutants ΔFsBmh1/sFsBmh2 and ΔFsBmh2/sFsBmh1, over-expression mutants O-FsBmh1 and O-FsBmh2, and complementation mutants C-ΔFsBmh1 and C-ΔFsBmh2, were stored in 25 % (v/v) glycerin at −80°C. For phenotypic assays, all strains were grown on potato dextrose agar medium (PDA) at 28°C for 7 days [22].

### Generation of mutant strains

To knockout *FsBmh1* and *FsBmh2*, the deletion fragments were amplified using fusion PCR with primer pairs FsBmh1-1F/R, FsBmh2-1F/R, FsBmh1-2F/R, FsBmh2-2F/R, Hph-F/R and G418-F/R (S1 Table). The hygromycin phosphotransferase gene (*hph*) and neo gene (*NeoR*) were amplified and used as the selection marker [23]. The resulting fragments were transformed individually into protoplasts of the wild type strain CNO-1 using a polyethylene glycol (PEG)-mediated transformation method to produce ΔFsBmh1 and FsBmh2 mutants [24]. To generate complementation strain, a 4093 bp and 4100bp fragments of the full length *FsBmh1* and *FsBmh2* including promoter sequences were amplified using primer pairs C-FsBmh1-F/R and C-FsBmh2-F/R (S1 Table). These fragments were cloned into the pCPXG418 or pCPXHY2 vector using pEASY-Basic Seamless Cloning and Assembly Kit (Transgen, Beijing, China), and then transformed into protoplasts of ΔFsBmh1 or ΔFsBmh2 mutant. The silenced strains were constructed using RNA interference method to silence *FsBmh1* in ΔFsBmh2 background and to silence *FsBmh2* in ΔFsBmh1 background. The 520 bp and 519 bp interference fragments for *FsBmh1* and *FsBmh2* (S4 Fig) were amplified from the genomic DNA of the wild-type strain with primers listed in Table S1 and the amplicons were cloned into the pCPXG418 or pCPXHY2 vector using pEASY-Basic seamless Cloning and Assembly Kit (Transgen, Beijing, China). The constructs were then used to transform protoplasts of ΔFsBmh2 or ΔFsBmh1 [25]. The overexpression strains were generated by transforming the protoplasts of the wild type strain CNO-1 with the *FsBmh1* or *FsBmh2* gene cloned into the pCPXHY2 or pCPXG418 (S5 Fig). Transformants were selected on media supplemented with corresponding antibiotics.

### RNA extraction and quantitative real-time RT-PCR

Total RNA of each strain was extracted with the RNA extraction kit (Takara biomedical technology, Beijing, China) following the manufacturer’s instruction. First-strand cDNA was synthesized with the FastQuant RT Kit (Takara biomedical technology, Beijing, China) following the manufacturer’s instructions. For quantitative real-time PCR, transcripts of the target genes were amplified with the corresponding primer pairs (S2 Table) and quantified with SuperReal PreMix Plus (SYBR Green) (Takara biomedical technology, Beijing, China), using the 18S rRNA was used as control. The relative quantification of the transcripts was calculated by the 2*^−^*^ΔΔCT^ method [26]. All qPCR assays were conducted with samples from three biological replications.

### Microscopy

Hyphae were stained with calcofluor white (CFW) [27] and viewed under an Olympus DP70 microscope.

### Pathogenicity assays

The sugarcane seedlings of susceptible cultivar Zhongzhe 9 (ZZ9) were grown in the greenhouse till 5 leaves stage before being used for inoculation. *F. sacchari* conidia were harvested from 7-days-old cultures grown on PDA plates and adjusted to 1×10^4^ conidia/ml with sterile distilled water. For inoculation, a volume of 500 *µ*l spore suspension was injected into sugarcane stem around the tip meristem using a sterile 22-gauge needle. Twenty-one days after inoculation (dpi), the disease symptom of the seedlings was evaluated. The disease severity index (DSI) were calculated by using a symptom severity scale ranging from 0 to 5 (S3 Table). The DSI was calculated using the formula DSI= 100× (score/ 5N), in which N is the number of sugarcane seedlings observed (N=100). A one-way ANOVA statistical analysis was performed using SPSS 23.0 software. Three replicates were performed for each assay.

### Transcriptomic analysis

Strains of interest were cultured in potato dextrose broth medium (PDB) at 28°C, 200 rpm for 3 days. Hyphae were harvested by filtration and washed three times with sterile distilled water. Total RNA was extracted from the tissue using TRIzol^®^ Reagent according the manufacturer’s instructions (Invitrogen) and genomic DNA was removed using DNase I (TaKaRa). The transcriptome libraries were constructed by using the TruSeqTM RNA sample preparation Kit from Illumina (San Diego, CA) and was sequenced using the Illumina NovaSeq 6000 platform at Shanghai Majorbio Bio-pharm Technology Co., Ltd. (Shanghai, China). Raw reads were filtrated to generate clean data using fastp v0.19.5. Clean reads were mapped to the genome of *F. sacchari* (gx1CNO-1) to obtain mapped data using TopHat2 v2.1.1. To identify differentially expressed genes (DEGs) between two different samples/groups, the expression level of each gene was calculated according to the fragments per kilobases per million reads (FPKM) method. RSEM (http://deweylab.biostat.wisc.edu/rsem/) was used to quantify gene abundances. A minimum of twofold difference in expression in the paired libraries (*log*_2_ratio −1 or *≥*1) was used as a standard to judge each of the differential expressed genes (DEGs) at the false discovery rate at *P* = 0.05 or less [28]. All identified DEGs were functionally annotated using the non-redundant NCBI protein databases of *Fusarium spp*. KEGG pathway enrichment analysis (http://www.genome.jp/kegg/) were performed for all the DEGs at the threshold of corrected *P*-adjust *<*0.05 [29]. Transcript of phenotype-associated genes were verified using cDNA of transcriptome samples as template and the fungal 18S RNA gene as internal standard. Data were from three biological replications.

### Statistics

Statistics were performed using one-way ANOVA using SPSS statistical package version 23.0 (SPSS Inc., United States). S-D-K test was performed to make comparisons between treatments, using a probability level of *P<*0.05.

## Results

### 14-3-3 proteins are highly conserved in *Fusarium* species

Two distinct genes (*FsBmh1* and *FsBmh2*) both encoding 14-3-3 proteins were identified in *F. sacchari* CNO1 genome using the *S. cerevisiae* Bmh1 and Bmh2 as queries. The cDNAs of FsBmh1 (FVER 07211) and FsBmh2 (FVER 01000) were 807 bp and 849 bp encoding 268 and 276 amino acids, respectively. These proteins have a typical 14-3-3 protein superfamily functional domain and share 69.6 % identity and 83.4 % similarity at amino acid level. Of interest were the facts that the amino acid homology of Bmh within the *Fusarium* genus (*F. sacchari*, *F. graminearum*, *F. fujikuroi*, and *F. oxysporum*) were 91.4 % to 97 %, while the homology between fungal genus was much lower, e.g., *S. cerevisiae* (62.4 %) and *C. albicans* (73.1 %) (S1A Fig). Phylogenetic analysis showed that Bmh1 and Bmh2 clustered separately among fungal species (S1B Fig).

### In vivo and in vitro expression patterns of *FsBmh1* and *FsBmh2* are discrepant

Sugarcane seedlings were inoculated with the spore suspension by infection. Hyphae development was observed to invade the plant at 24 hpi; hyphal branching, plant cell disintegration and necrotic spots on the surface of sugarcane leaves were observed at 48 hpi; and severe tissue necrosis, hyphae spread in the infected cells and new spores were found at 96 hpi (Fig 1A and Fig 1B). Corresponding to these stages, *FsBmh1* transcript level elevated by 2-3 folds in 24 and 48 h, but sharply lifted to 41 folds in 96 h, as compared with 0 h; *FsBmh2* transcript level did not changed in the first 24 h, but upregulated by about 2 folds at 48 h, and 15 folds at 96 h (Fig 1C). Although both *FsBmh1* and *FsBmh2* transcriptionally responded in colonization to the host plant, *FsBmh1* responded earlier than *FsBmh2* in planta.

**Fig 1.**
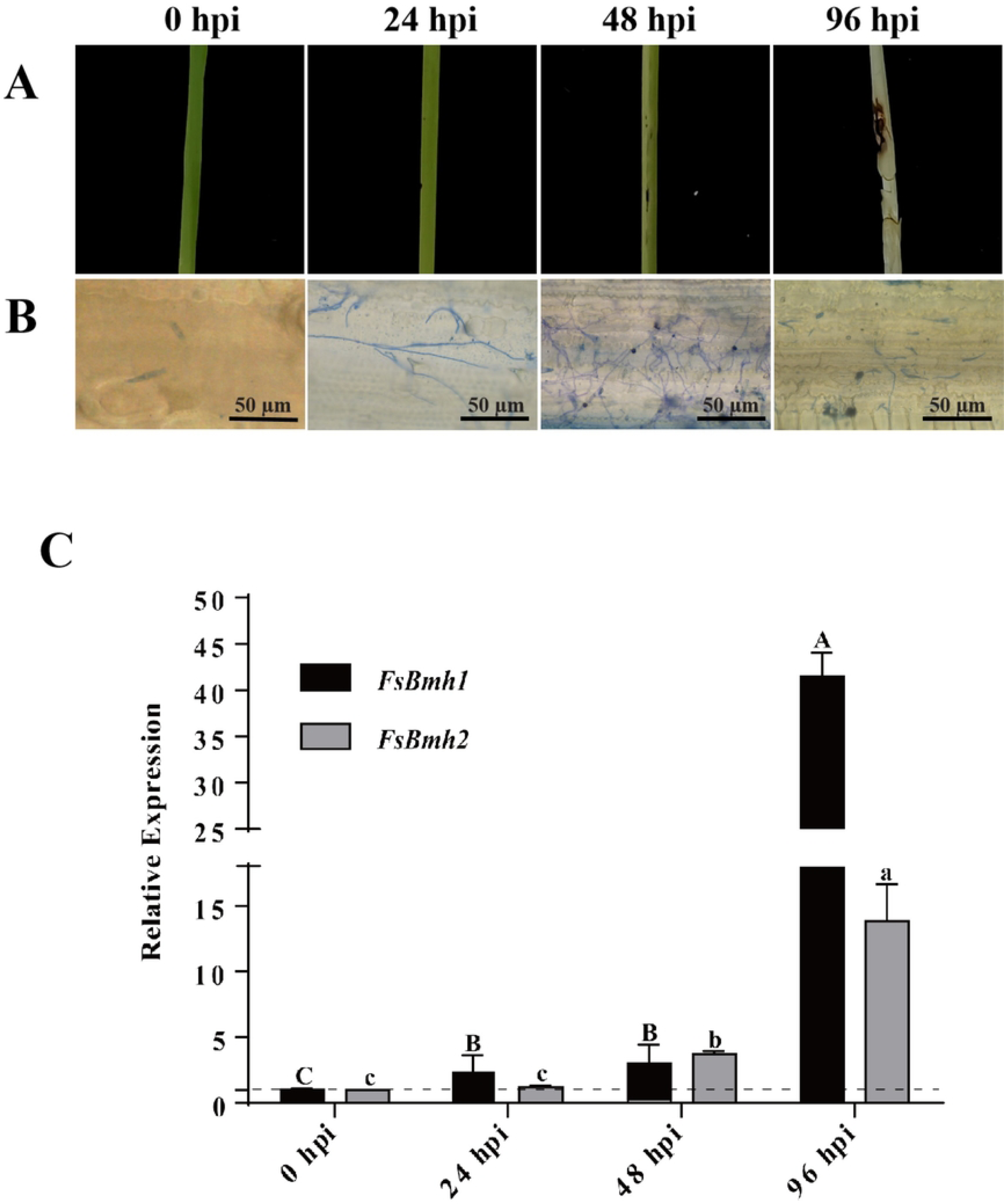
Expression patterns of *FsBmh1* and *FsBmh2* at different developmental stages in vivo. (A) The sugarcane seedlings were inoculated with CNO-1, and representative seedlings were pohotographed at 24 hpi, 48 hpi and 96 hpi. (B) Sections were stained by Aniline Blue and observed under microscope. (C) Relative expression of *FsBmh1* and *FsBmh2* in infection stages. The mean and standard deviation were derived from three independent biological replicates, each of which contained three technical repeats. The expression of *FsBmh1* and *FsBmh2* transcripts was measured by quantitative real-time RT-PCR (2*^−^*^ΔΔCT^ method). The relative transcript level was calibrated using 18S rRNA as reference, and the transcript levels of WT were set to a value of one. Different letters indicate significant differences of each strain (*P<*0.05).

To investigate the developmental time course of *F. sacchari* and transcription dynamics of *FsBmh* in saprophytic condition, we inoculated spores of *F. sacchari* in the PDB medium. It was found that initial hyphal growth was observed at 10 h, branching at 14 h, and new spore formation at 22 h (Fig 2A). Quantification of transcript levels revealed that *FsBmh1* was upregulated by 3.3 folds at 10 h, then declined (1.4 folds at 14 h and 0.8 fold at 22 h); however, *FsBmh2* was upregulated by 4.3 folds at 10 h and maintained at about the same level onward (Fig 2B). By comparison of the transcription profiles, it was clear that there was a discrepancy in transcription dynamics between *FsBmh1* and *FsBmh2* both in vivo and in vitro.

**Fig 2.**
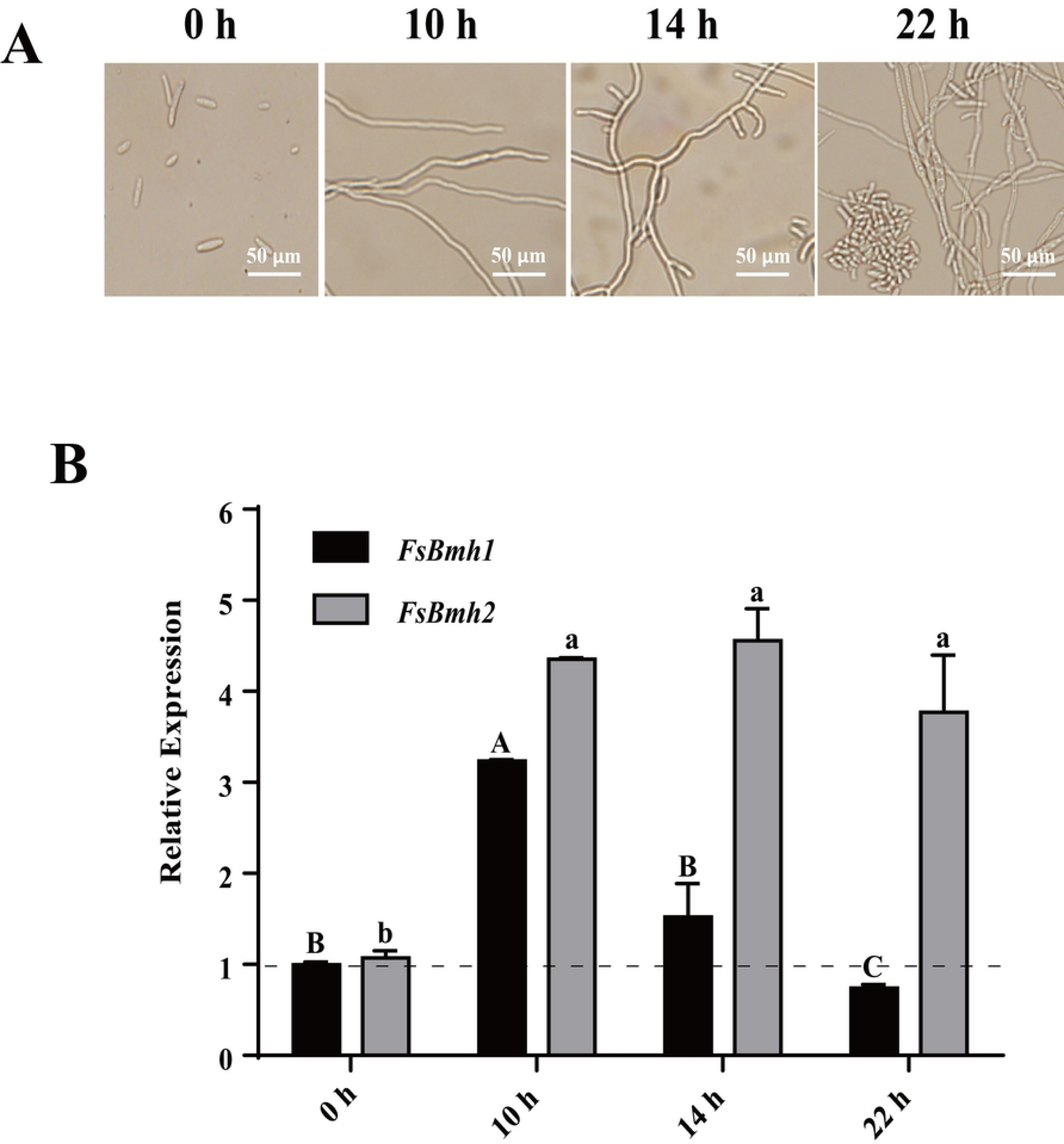
Expression patterns of *FsBmh1* and *FsBmh2* at different developmental stages in vitro. (A) The spore suspension of CNO-1 were inoculated in PDB medium, and representative pictures were pohotographed at 10 h, 14 h and 22 h by microscope. (B) Relative expression of *FsBmh1* and *FsBmh2* in saprophytic stages. The mean and standard deviation were derived from three independent biological replicates, each of which contained three technical repeats. The expression of *FsBmh1* and *FsBmh2* transcripts was measured by quantitative real-time RT-PCR (2*^−^*^ΔΔCT^ method). The relative transcript level was calibrated using 18S rRNA as reference, and the transcript levels of WT were set to a value of one. Different letters indicate significant differences of each strain (*P<*0.05).

### The expression of *FsBmh1* and *FsBmh2* are mutually compensated in cells

To elucidate the distinct functions of two 14-3-3 proteins, we generated gene deletion mutants ΔFsBmh1 and ΔFsBmh2 by homologous recombination (S2 Fig and S3 Fig). Intriguingly, deletion of *FsBmh1* resulted in the elevated expression of *FsBmh2* (2.6-fold higher) and deletion of *FsBmh2* resulted in the elevated expression of *FsBmh1* (1.8-fold higher). However, reintroduction of a wild copy of *FsBmh1* or *FsBmh2* into the corresponding mutants (C-ΔFsBmh1 and C-ΔFsBmh2), abolished this expression compensation (Fig 3).

**Fig 3.**
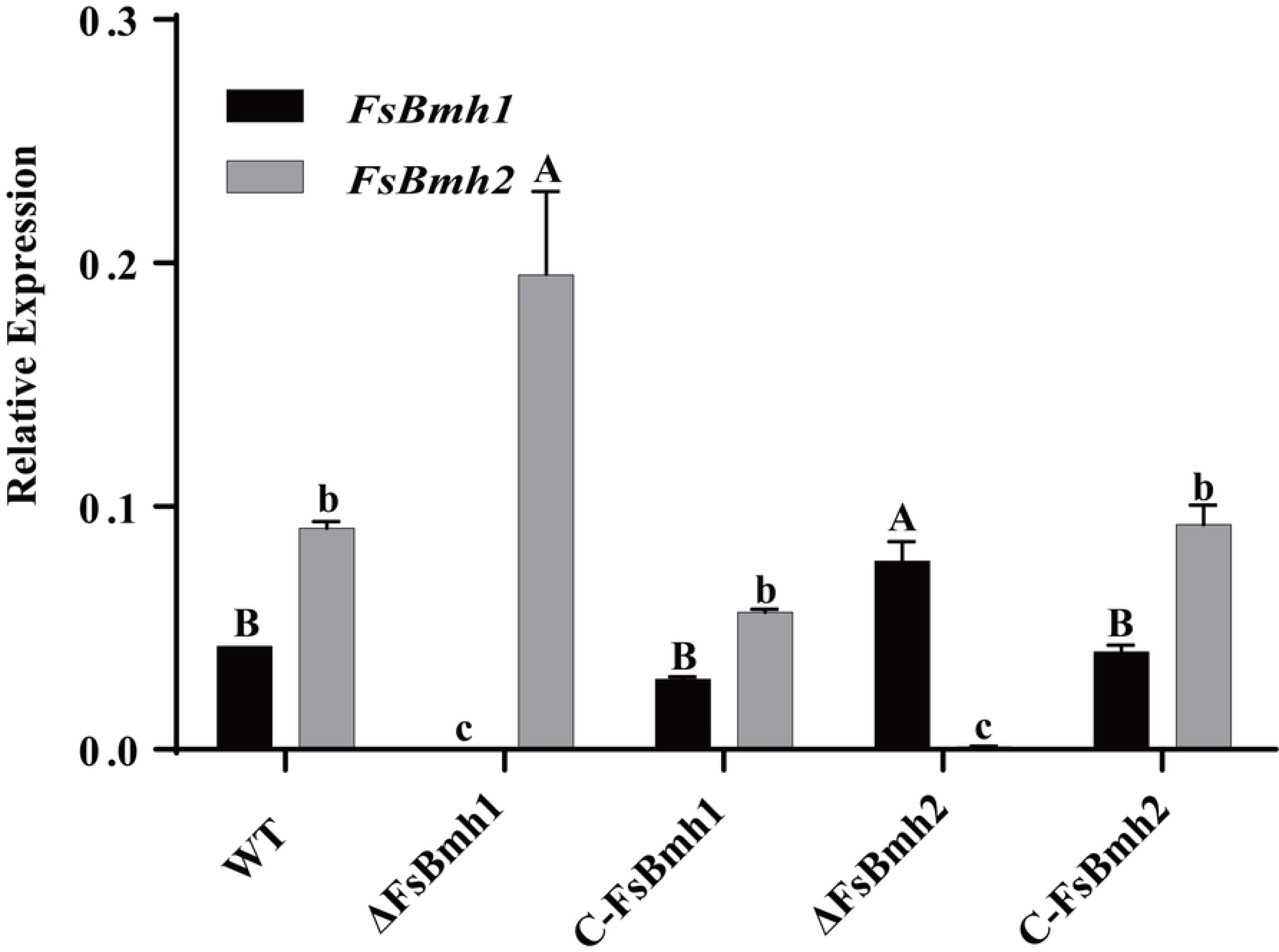
Validation of *FsBmh1* and *FsBmh2* expression using qRT-PCR in deletion and complementation mutants. Relative expression of *FsBmh*-genes in *FsBmh*-deletion and complementation mutants were determined by qRT-PCR. The mean and standard deviation were derived from three independent biological replicates, each of which contained three technical repeats. Relative expression of *FsBmh1* and *FsBmh2* were measured using 18S rRNA as reference, whose level was set as 1.0. Different letters indicate significant differences of each strain (*P<*0.05).

### *FsBmh1* and *FsBmh2* contribute to fungal phenotypes in varied ways

Compared with the wild type strain, ΔFsBmh1 and ΔFsBmh2 lost the light brown pigment and grew slowly by about 10% and 30%. Of particular note, deletion of *FsBmh1* resulted in longer but thinner microspores, and deletion of *FsBmh2* resulted in an increasing number of branching, significantly shorter mycelial cell and smaller microspores (Fig 4A and Table 1 and Table 2). These results demonstrated that both *FsBmh1* and *FsBmh2* contribute to regulate the cell architecture but in different ways in *F. sacchari*. Quantification of microconidia revealed that both *FsBmh1* and *FsBmh2* contributed to sporulation, but the impact of *FsBmh2* was significantly greater than *FsBmh1*, e.g. deletion of *FsBmh1* reduced spore yield by 43%., but deletion of *FsBmh2* reduced spore yield by 80%. (Fig 4B). Moreover, spores from the ΔFsBmh2 germinated at a lower rate of 67% as compared with the wild type, while ΔFsBmh1 did not seem to affect the germination rate (Fig 4C).

**Fig 4.**
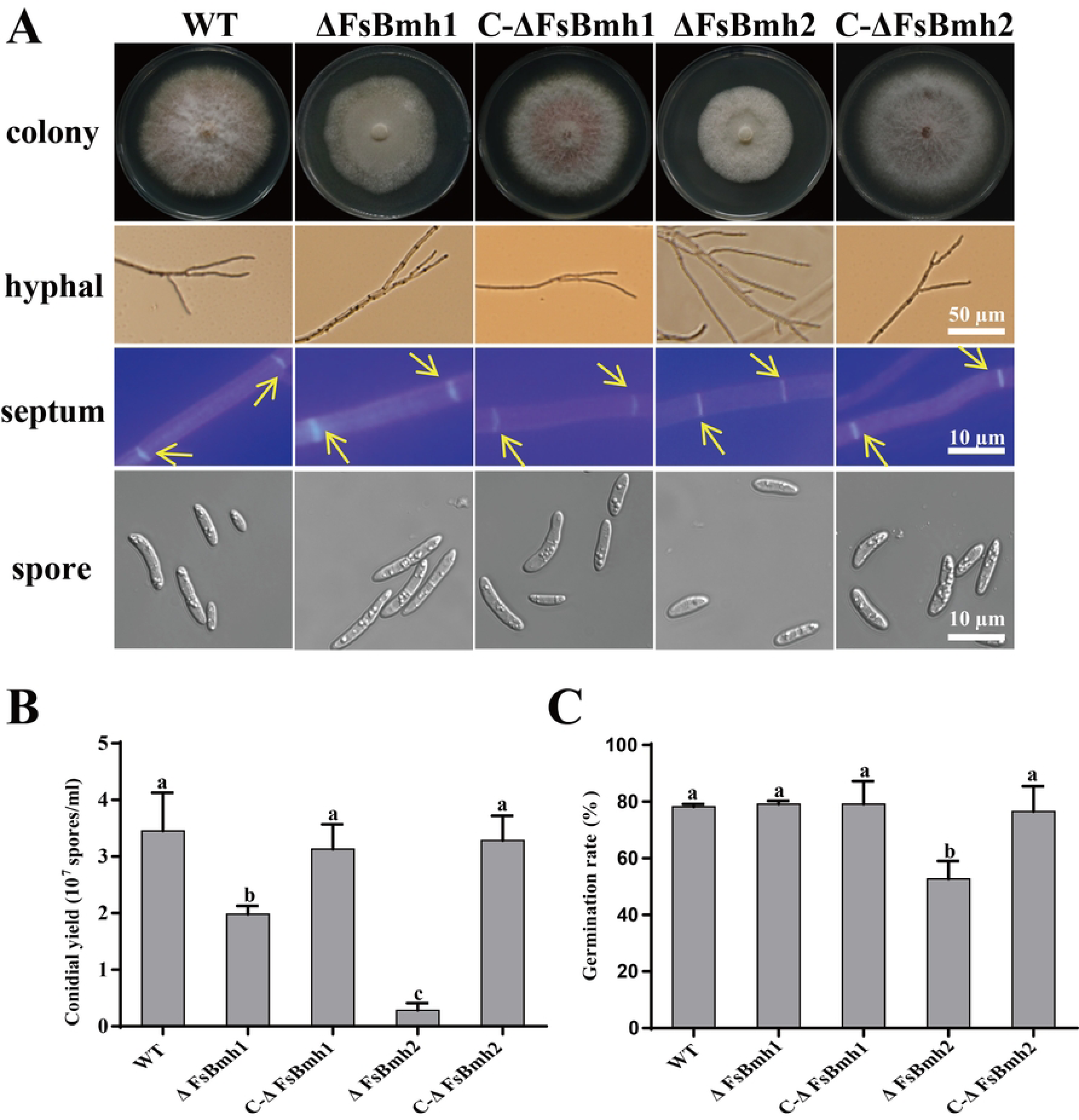
Phenotypes of *FsBmh*-deletion mutants and complementation mutants. (A) Hyphal and conidial phenotypes of strains. Colony were photographed on PDA plates for 7 days. Strains were inoculated on PDA for 3 days and recorded hyphal branching using microscope, scale bar = 50 *µ*m. Hyphal separation was observed by staining with CFW and photographed by microscope, scale bar = 10 *µ*m. The separations were marked by yellow arrows. Spore characteristics were photographed by microscope, scale bar = 10 *µ*m. (B) Statistics of conidial yield of strains. The number of Conidial harvested from PDA plates for 7 days. (C) Conidial germination rate of strains were recorded after 6 h of incubation in PDB medium at 28°C,150 rpm. The mean and standard deviation were calculated from three replicates. Different letters indicate significant differences of each strain (*P<*0.05)

**Table 1.** Cell length measurement of mutant strains.

| strain | Hyphae with different septa(%) | | | | Hyphal length<br>with three septa ($\mu\text{m}$) | Cell length ($\mu\text{m}$) |
| --- | --- | --- | --- | --- | --- | --- |
|  | 1 | 2 | 3 | 4 |  |  |
| WT | 25 | 19 | 35 | 21 | 82.6 ± 10.57<sup>a</sup> | 20.0 ± 3.86<sup>a</sup> |
| $\Delta\text{FsBmh1}$ | 31 | 25 | 27 | 17 | 79.9 ± 11.37<sup>a</sup> | 19.9 ± 0.86<sup>a</sup> |
| O-FsBmh1 | 27 | 17 | 26 | 30 | 88.5 ± 17.98<sup>a</sup> | 24.5 ± 2.12<sup>a</sup> |
| $\Delta\text{FsBmh1/sFsBmh2}$ | 25 | 21 | 32 | 22 | 44.9 ± 8.33<sup>b</sup> | 12.6 ± 1.08<sup>b</sup> |
| C-$\Delta\text{FsBmh1}$ | 21 | 23 | 36 | 20 | 82.4 ± 7.12<sup>a</sup> | 20.8 ± 3.34<sup>a</sup> |
| $\Delta\text{FsBmh2}$ | 31 | 23 | 31 | 15 | 39.8 ± 7.78<sup>b</sup> | 11.9 ± 1.13<sup>b</sup> |
| O-FsBmh2 | 25 | 17 | 32 | 26 | 94.8 ± 14.73<sup>a</sup> | 22.1 ± 0.24<sup>a</sup> |
| $\Delta\text{FsBmh2/sFsBmh1}$ | 19 | 23 | 39 | 19 | 35.2 ± 7.43<sup>b</sup> | 9.9 ± 1.03<sup>b</sup> |
| C-$\Delta\text{FsBmh2}$ | 19 | 19 | 31 | 31 | 86.8 ± 11.92<sup>a</sup> | 22.1 ± 2.31<sup>a</sup> |

Percentages of hyphal with different septa (n = 100) in wild-type and all mutant strains were determined after 8 h of incubation in PDB medium, at 28 °C,150 rpm. Hyphal length with three septa and cell length were measured by Image J. The mean and standard deviation were derived from one-hundred replicates. Different letters indicate significant differences of each strain ( $P<0.05$ ).

**Table 2.** Spore length and width measurement of all strains inoculated on PDA.

| Strain | Spore length ($\mu\text{m}$) | Spore width ($\mu\text{m}$) |
| --- | --- | --- |
| WT | 13.63±0.54<sup>b</sup> | 3.38±0.21<sup>a</sup> |
| $\Delta\text{FsBmh1}$ | 17.92±1.02<sup>a</sup> | 2.35±0.27<sup>b</sup> |
| O-FsBmh1 | 13.92±1.87<sup>b</sup> | 3.35±0.20<sup>a</sup> |
| $\Delta\text{FsBmh1/sFsBmh2}$ | 16.60±1.70<sup>a</sup> | 3.38±0.12<sup>a</sup> |
| C-$\Delta\text{FsBmh1}$ | 14.05±0.26<sup>b</sup> | 3.44±0.16<sup>a</sup> |
| $\Delta\text{FsBmh2}$ | 10.85±0.70<sup>c</sup> | 3.55±0.15<sup>a</sup> |
| O-FsBmh2 | 13.99±1.26<sup>b</sup> | 3.31±0.17<sup>a</sup> |
| $\Delta\text{FsBmh2/sFsBmh1}$ | 10.79±1.26<sup>c</sup> | 2.54±0.38<sup>b</sup> |
| C-$\Delta\text{FsBmh2}$ | 14.04±0.61<sup>b</sup> | 3.33±0.14<sup>a</sup> |

Spores were harvested after incubation for 7 days on PDA medium. Spore length and width were measured by Image J. The mean and standard deviation were derived from one-hundred replicates. Different letters indicate significant differences of each strain ( $P<0.05$ ).

### *FsBmh1* and *FsBmh2* interactively regulate spore morphology in *F. sacchari*

We attempted to knockout both *FsBmh1* and *FsBmh2* in the same strain but failed to recover any such mutants. This may suggest that Spontaneous deletion of *FsBmh1* and *FsBmh2* lead to lethal phenotype in *F. sacchari*. Alternatively, we constructed a *FsBmh2*-silienced strain (15.3% of the *FsBmh2* expression) in ΔFsBmh1 (ΔFsBmh1/sFsBmh2) and *FsBmh1*-silienced (13.9% of the *FsBmh1* expression) in ΔFsBmh2 (ΔFsBmh2/sFsBmh1) (Fig 5B). RNAi of *FsBmh* only showed a marginal inhibition effect on the colonial growth, i.e. 8% on theΔFsBmh1/sFsBmh2 as compared with ΔFsBmh1, and 5% on ΔFsBmh2/sFsBmh1 as compared with ΔFsBmh2. However, more branches and shorter mycelial cell were observed in ΔFsBmh1/sFsBmh2, similar to the characteristics of the ΔFsBmh2 (Fig 6 and Table 1). Unexpectedly, larger and longer macrospores were observed in ΔFsBmh1 and ΔFsBmh1/sFsBmh2 (Fig 7 and Table 3), suggesting that production of macrospores was also regulated by *FsBmh1* and *FsBmh2*. Of interest was the observation that reduction in conidiation could be compounded by RNAi of *FsBmh2* on ΔFsBmh1 (ΔFsBmh1/sFsBmh2) but this accumulated effect was not obvious in ΔFsBmh2/sFsBmh1 (Fig 6B). However, accumulated adverse effect on germination was seen in ΔFsBmh2/sFsBmh1 for which spore germination rate of 37% was significantly lower than the 52% forΔFsBmh2, while ΔFsBmh1 and ΔFsBmh1/sFsBmh2 were at basically the same germination rate as the wild type (Fig 6C).

**Fig 5.**
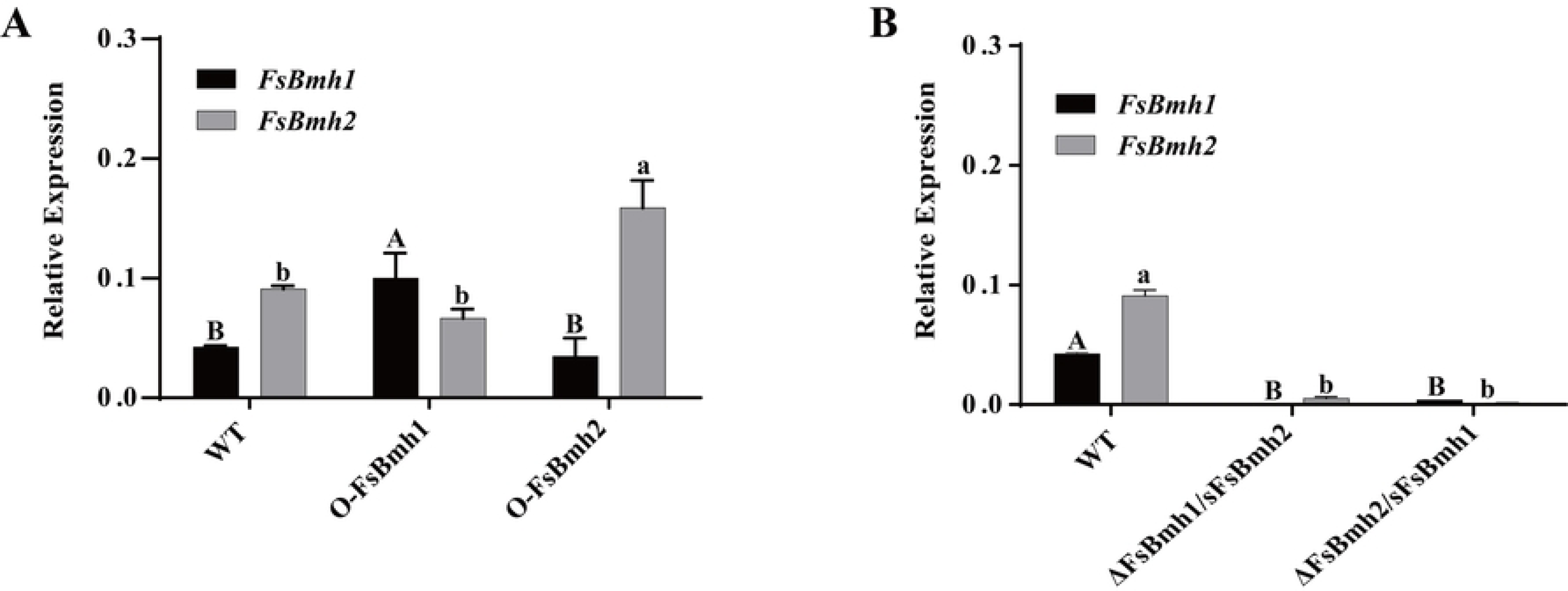
qRT-PCR validation of *FsBmh1* and *FsBmh2* expression in *FsBmh*-silenced mutants and overexpression strains. (A) Relative expression of *FsBmh* genes in O-FsBmh1 and O-FsBmh2 mutants were determined by qRT-PCR. (B) Relative expression of *FsBmh1* and *FsBmh2* genes in ΔFsBmh1/sFsBmh2 and ΔFsBmh2/sFsBmh1 mutants were determined by qRT-PCR. The mean and standard deviation were derived from three independent biological replicates, each of which contained three technical repeats. Relative expression of *FsBmh1* and *FsBmh2* were measured using 18S rRNA as reference, whose expression level was set as 1.0. Different letters indicate significant differences of each strain (*P<*0.05).

**Fig 6.**
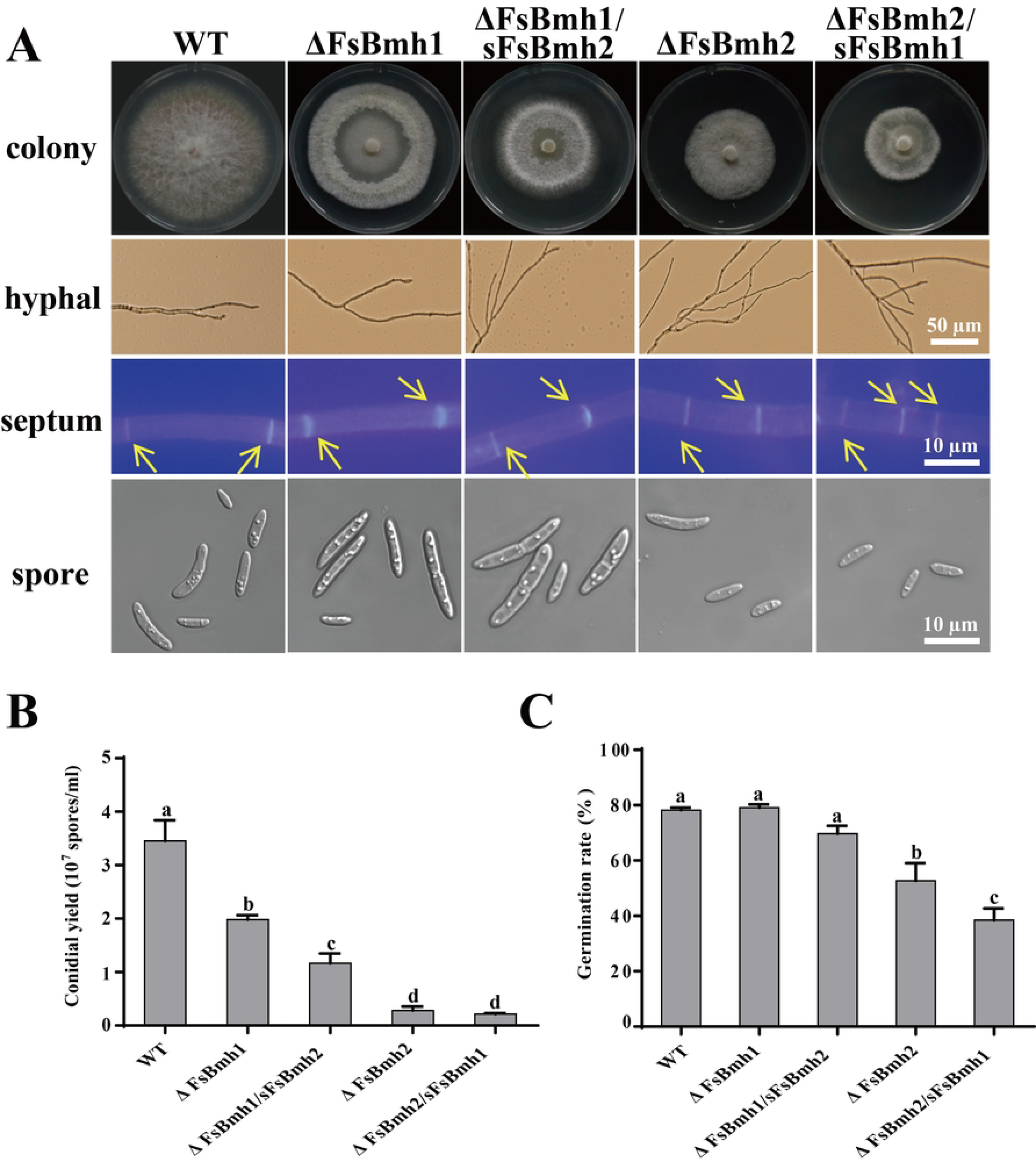
Phenotypes of *FsBmh*-deletion mutants and *FsBmh*-silenced mutants. (A) Hyphal and conidial phenotypes of strains. Colony were photographed on PDA plates for 7 days. Strains were inoculated on PDA for 3 days and recorded hyphal branching using microscope, scale bar = 50 *µ*m. Hyphal separation was observed by staining with CFW and photographed by microscope, scale bar = 10 *µ*m. The separations were marked by yellow arrows. Spore characteristics were photographed by microscope, scale bar = 10 *µ*m. (B) Statistics of conidial yield of strains. The number of Conidial harvested from PDA plates for 7 days. (C) Conidial germination rate of strains were recorded after 6 h of incubation in PDB medium at 28°C,150 rpm. The mean and standard deviation were calculated from three replicates. Different letters indicate significant differences of each strain (*P<*0.05).

**Fig 7.**
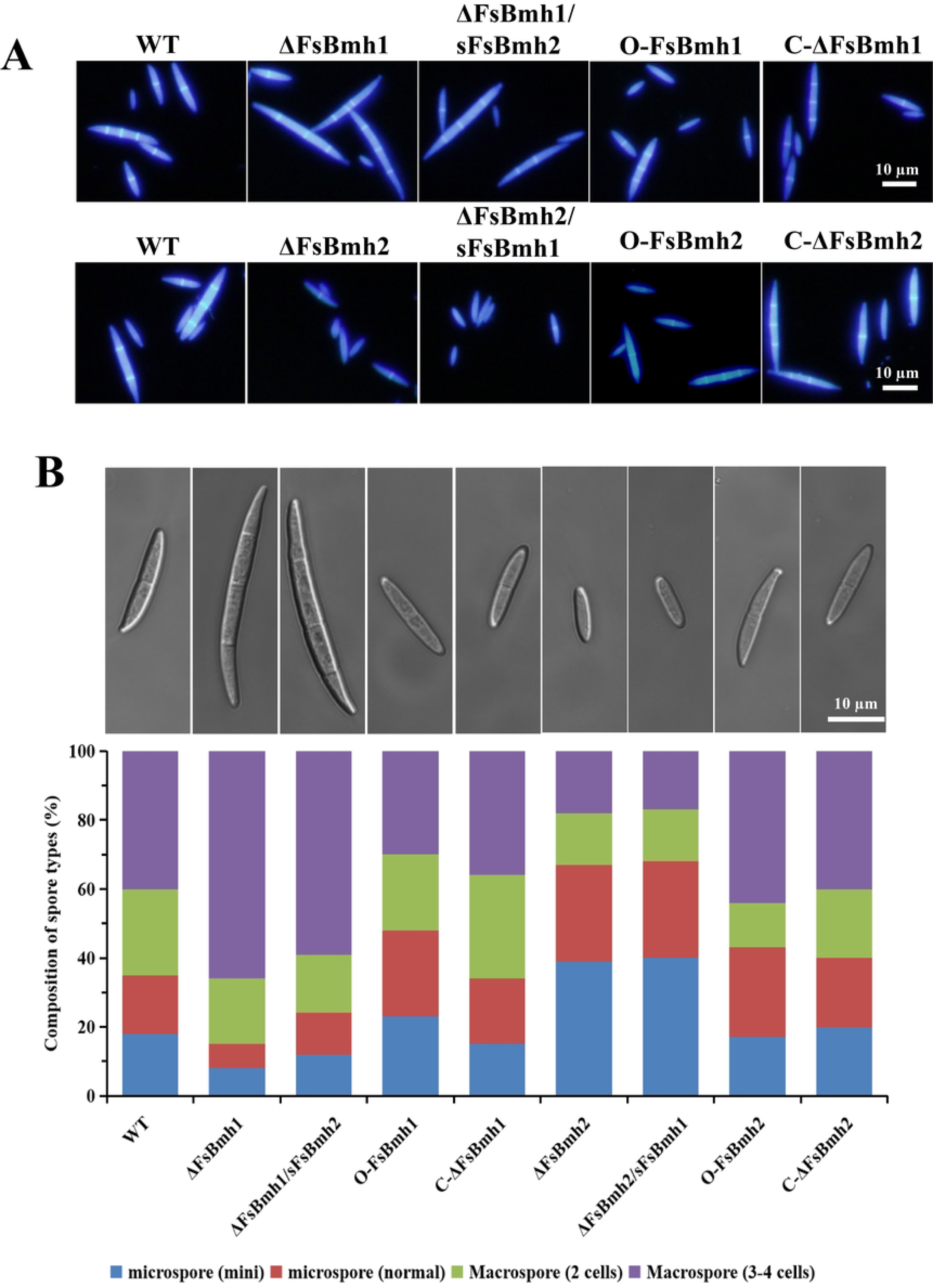
FsBmh1 and *FsBmh2* contribute to macrospore production and morphology. (A) Macrospores of CNO-1 and all mutant strains were induced by carnation leaves. Spores including microspore (mini), microspore (normal), macrospore (2 cells) and macrospore (3-4 cells) were harvested and stained by calcofluor white and examined by fluorescent microscope. (B) Spore types of CNO-1 and all mutant strains were counted (n = 100). The images were taken using differential-interference contrast microscope and represent the spore characteristics that occur frequently in each strain.

**Table 3.** Spore length and width measurement of all strains inoculated on PDA.

| strain | Spore length ($\mu\text{m}$) | | | | Spore width ($\mu\text{m}$) | | | |
| --- | --- | --- | --- | --- | --- | --- | --- | --- |
|  | Microspore (mini) | Microspore (normal) | Macrospore (2 cells) | Macrospore (3-4 cells) | Microspore (mini) | Microspore (normal) | Macrospore (2 cells) | Macrospore (3-4 cells) |
| WT | 8.32±1.52a | 14.96±2.29a | 22.10±1.69ab | 26.90±2.96b | 1.68±0.45a | 1.98±0.43ab | 2.22±0.41ab | 2.46±0.51ab |
| $\Delta\text{FsBmh1}$ | 8.33±1.23a | 14.64±2.96a | 24.55±3.54ab | 34.91±3.11a | 1.70±0.41a | 1.81±0.25b | 1.94±0.30b | 2.10±0.25b |
| $\Delta\text{FsBmh1/sFsBmh2}$ | 8.21±1.93a | 14.12±2.68a | 25.06±3.08a | 32.44±2.87a | 1.74±0.45a | 1.94±0.18ab | 2.27±0.46ab | 2.36±0.53ab |
| O-FsBmh1 | 8.58±1.24a | 14.83±1.80a | 22.75±1.65ab | 25.17±1.83b | 1.85±0.31a | 1.96±0.14ab | 2.29±0.45ab | 2.44±0.48ab |
| C-$\Delta\text{FsBmh1}$ | 8.41±2.90a | 15.25±2.99a | 22.83±3.04ab | 25.17±2.23b | 1.75±0.45a | 2.08±0.29ab | 2.20±0.45ab | 2.42±0.51ab |
| $\Delta\text{FsBmh2}$ | 6.89±1.71ab | 13.61±2.67a | 21.60±2.19b | 25.06±1.67b | 1.94±0.24a | 2.33±0.49ab | 2.60±0.55a | 2.77±0.42a |
| $\Delta\text{FsBmh2/sFsBmh1}$ | 6.10±1.19b | 13.44±2.00a | 21.87±2.42b | 24.75±1.28b | 1.85±0.34a | 2.11±0.33ab | 2.31±0.46ab | 2.50±0.53ab |
| O-FsBmh2 | 8.73±2.24a | 15.18±1.78a | 22.73±1.73ab | 25.36±2.42b | 1.75±0.40a | 2.09±0.30ab | 2.23±0.41ab | 2.42±0.43ab |
| C-$\Delta\text{FsBmh2}$ | 8.54±3.07a | 15.09±1.87a | 22.54±2.16<sup>ab</sup> | 25.18±2.44b | 1.81±0.34a | 2.04±0.35b | 2.25±0.43ab | 2.42±0.48ab |

Spores were harvested after incubation for 7 days on carnation leaf-piece agar medium. Spore length and width were measured by Image J. The mean and standard deviation were derived from one-hundred replicates. Different letters indicate significant differences of each strain ( $P<0.05$ ).

### Overexpression *FsBmh* does not affect the phenotype of *F.sacchari*

We constructed *FsBmh1* - and *FsBmh2*-overexpression strains with transcripts elevated by 135.6% and 66.4%, respectively (Fig 5A). No significant difference in phenotypes, e.g. hyphal growth, hyphal branching, cell length, sporulation, spore morphology and germination rate between the O-FsBmh and the wild-type strain was observed (Fig 8).

**Fig 8.**
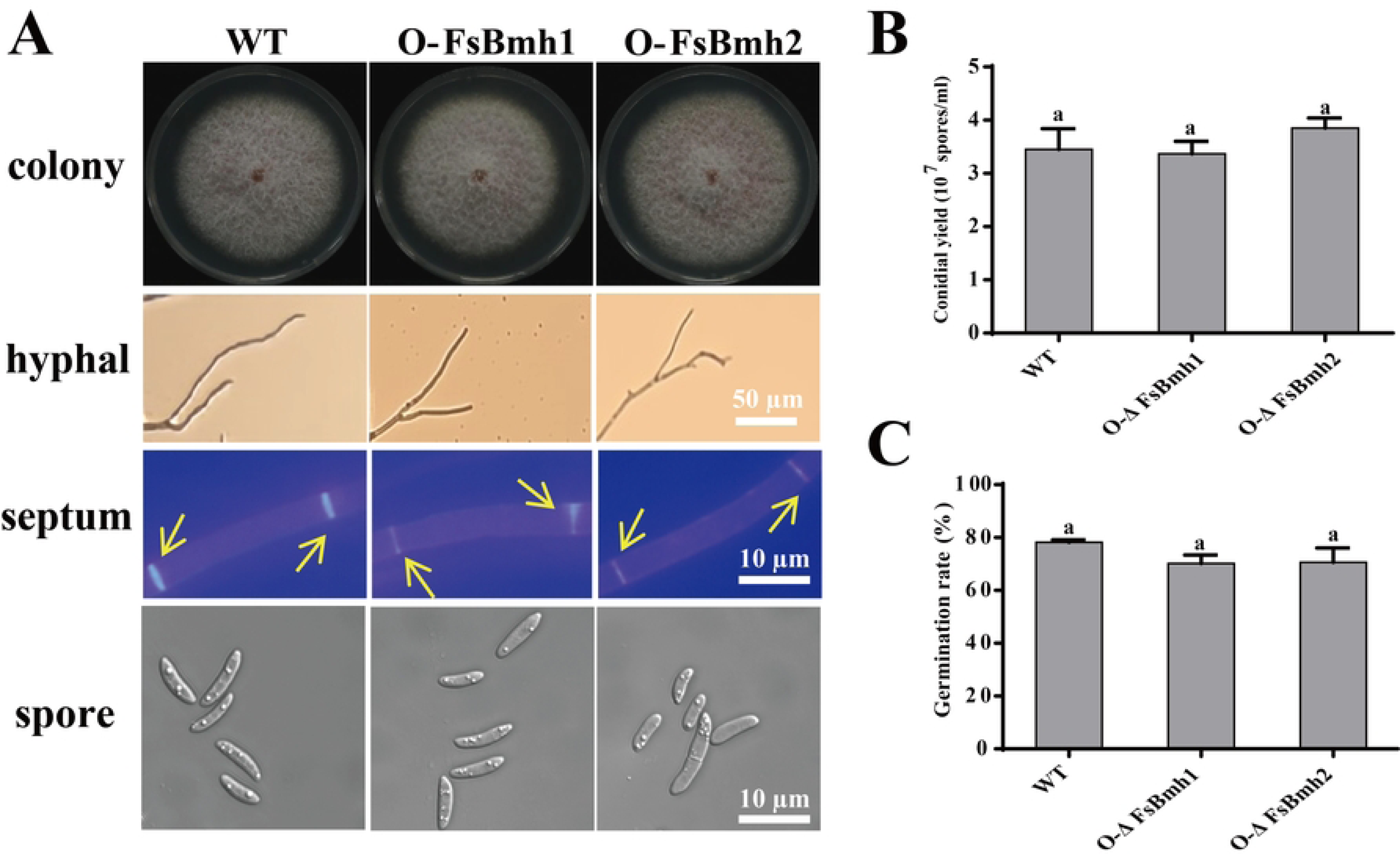
Phenotypes of - overexpression mutants. (A) Hyphal and conidial phenotypes of strains. Colony were photographed on PDA plates for 7 days. Strains were inoculated on PDA for 3 days and recorded hyphal branching using microscope, scale bar = 50 *µ*m. Hyphal separation was observed by staining with CFW and photographed by microscope, scale bar = 10 *µ*m. The separations were marked by yellow arrows. Spore characteristics were photographed by microscope, scale bar = 10 *µ*m. (B) Statistics of conidial yield of strains. The number of Conidial harvested from PDA plates for 7 days. (C) Conidial germination rate of strains were recorded after 6 h of incubation in PDB medium at 28°C,150 rpm. The mean and standard deviation were calculated from three replicates. Different letters indicate significant differences of each strain (*P<*0.05).

### *FsBmh1* and *FsBmh2* are both required for virulence to the sugarcane

To test whether *FsBmh1* and *FsBmh2* influence the virulence in *F. sacchari*, we inoculated sugarcane plants with conidia (1×10^4^ conidia/ml). Typical PBD symptoms appeared on plants inoculated with the wild-type strains at 14 dpi, but significantly milder symptoms on plants inoculated with ΔFsBmh1, ΔFsBmh2, ΔFsBmh1/s FsBmh2, or ΔFsBmh2/sFsBmh1 at 21 dpi. The virulence of the *FsBmh2* could be fully restored by re-introduction of a copy of the wild-type *FsBmh2*, while only about 60% of the virulence could be restored by re-introduction of a copy of the wild-type *FsBmh1* into the deletion of *FsBmh1* mutant, as judged by disease severity index. A significant increase in virulence was found in the strain overexpressing *FsBmh1*, but not in the *FsBmh2*-overexpressing strain (Fig 9).

**Fig 9.**
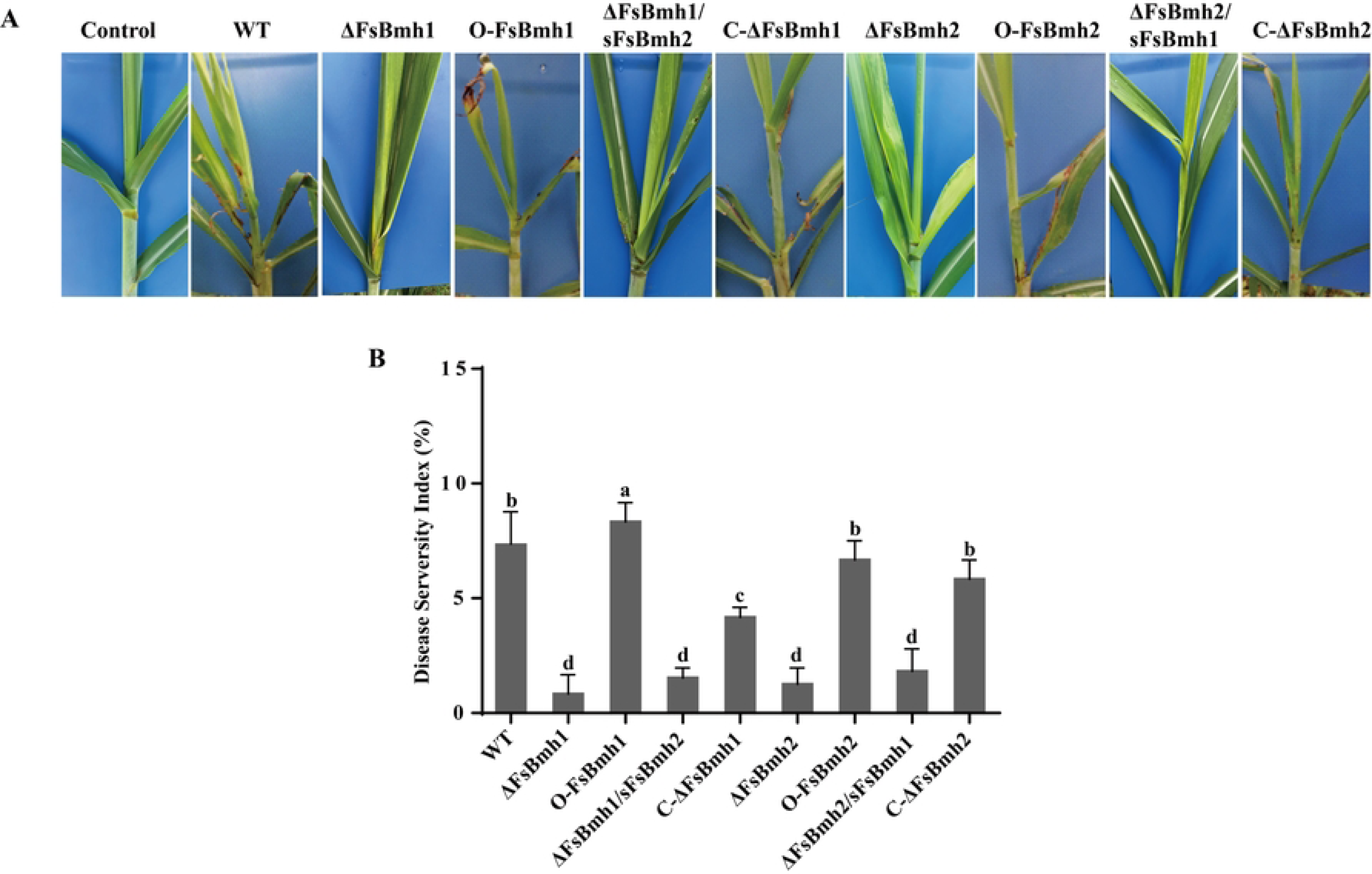
*FsBmh*-deletion mutants are defective in virulence. (A) Sugarcane seedlings were inoculated with the wide type and all mutant strains and photographed after 21dpi. (B) Disease severity index of all strains were determined using 100 seedlings per treatment at 21dpi. Different letters indicate significant differences of each strain (*P<*0.05).

### Transcriptomic basis for the functionality of *FsBmh1* and *FsBmh2*

To explore the mechanisms by which *FsBmh1* and *FsBmh2* regulate the fungal phenotypes, comparative transcriptomic analysis was performed on ΔFsBmh1, ΔFsBmh2 and WT. A total of 4296 differentially expressed genes (DEGs), 2059 up- and 2237 down-regulated, between ΔFsBmh1 and WT, and 4921 DEGs (2376 up- and 2542 down-regulated) between ΔFsBmh2 and WT were detected. As expected, ΔFsBmh1 and ΔFsBmh2 shared 3804 DEGs, including 1801 up-regulated genes and 2003 down-regulated genes (Fig 10A). Kyoto Encyclopedia of Genes and Genomes (KEGG) enrichment analysis (http://www.genome.jp/kegg/) showed that 265 up-regulated genes in ΔFsBmh1 and 308 up-regulated genes in ΔFsBmh2 were mainly enriched in ribosome biogenesis, RNA polymerase and RNA transport, mismatch repair and DNA replication. However, up-regulated DEGs of ΔFsBmh2 were specifically enriched in amino acid and propanoate metabolism pathways. On the other side, 184 and 224 down-regulated genes inΔFsBmh1 andΔFsBmh2 were mainly enriched in carbohydrate metabolism, amino acid metabolism, lipid metabolism and energy metabolism (Fig 10B).

**Fig 10.**
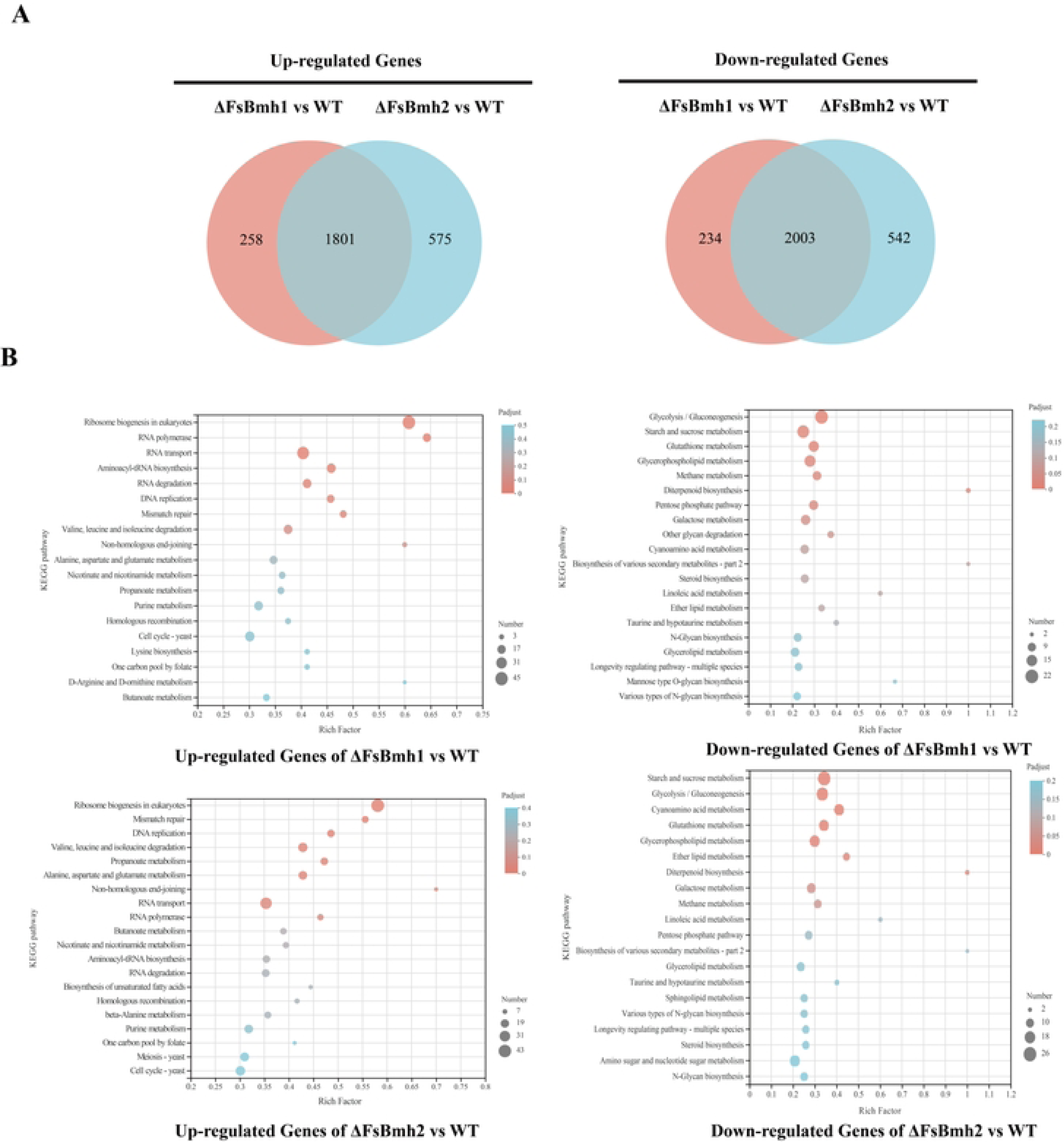
Distribution of DEGs (*≥* twofold) in ΔFsBmh1 and ΔFsBmh2 versus WT. (A) Venn diagrams showing the overlapped counts of the genes upregulated or downregulated in bothΔFsBmh1 and ΔFsBmh2 versus WT. (B) KEGG pathway enrichment of genes upregulated or downregulated in ΔFsBmh1 and ΔFsBmh2 versus WT.

A hierarchical clustering analysis of the DEGs further unveiled that the expression patterns of 981 genes that enriched in KEGG pathways (ΔFsBmh-versus WT) could be divided into 10 clusters (Fig 11A). We focused on the five representative clusters (cluster II, cluster III, cluster VI, cluster VIII, clusterX) as shown in Fig 11B. In cluster X and cluster II, 33 and 87 genes were down-regulated in all deletion strains vs WT and the expression level of ΔFsBmh2 was lower than ΔFsBmh1. Most of the genes were enriched in catalytic activity category by gene ontology (GO) enrichment, including four genes (FVER 13707, FVER 12121, FVER 07617, FVER 10685) encoding phosphatidylserine decarboxylase 2, ABC transporter, chitin synthase and S-(hydroxymethyl) glutathione dehydrogenase respectively, which is critical for virulence, sporulation, hyphal growth and transport of nutrients and efflux of antifungal drugs [30–33]. In cluster III, 182 genes were depressed at least two-fold in all deletion strains, but the expression level of ΔFsBmh1 were lower than ΔFsBmh2. These genes were enriched into phospholipid biosynthetic process, phosphorus metabolic process, organophosphate metabolic process and cellular lipid metabolic process. Phospholipase D (FVER 13005) was identified in all categories, it produces fused lipid phosphatidic acid, which acts as a second messenger regulates a variety of intracellular activities, including sporulation, mitosis, actin-cytoskeleton reorganization and virulence [34, 35]. Additionally, the gene encoding adenylate cyclase (FVER 07023) was appeared in cluster III. In cluster VIII, 12 genes were repressed in ΔFsBmh2 but up-regulated in ΔFsBmh1 vs WT. Most of genes were enriched in catalytic activity category, including two chitinase synthesis genes(FVER 00509, FVER 06234) and a gene (FVER 08232) encoding catalase. In cluster VI, 450 genes were up-regulated in all deletion strains and the expression of ΔFsBmh2 was higher thanΔFsBmh1. Most of genes were enriched in DNA repair, mismatch repair, DNA metabolic process and cell to response stress categories by gene ontology (GO) enrichment, such as encoding heat shock protein SSB1 (FVER 06187), the genes were up-regulated by at least two-fold and were reported to be involved in regulate cell signal transduction. There are two genes (FVER 08397,FVER 02730) were classified into DNA repair category, encoding DNA repair protein RAD50 and double-strand break repair protein MRE11, respectively. They are often present in the cell as a Rad50-Mre11-Xrs2 complex and play a central role in the reaction to DNA double-strand breaks [36]. Type 2A phosphatase (FVER 05113) a member of TOR signal pathway, may interact with FgPpg1 regulate hyphal development and virulence in *F. graminearum* [37]. Additionally, the gene encoding cell division control protein 45 (FVER 01583) were identified in this cluster, the expression were up-regulated by 4.9-fold in ΔFsBmh2. CDC45 has been shown to be associated with the process of DNA replication initiation and is a negative regulator of cell proliferation [38]. Relative expression of these five genes was validated by qRT-PCR, consistent with transcriptomic data (Fig12A).

**Fig 11.**
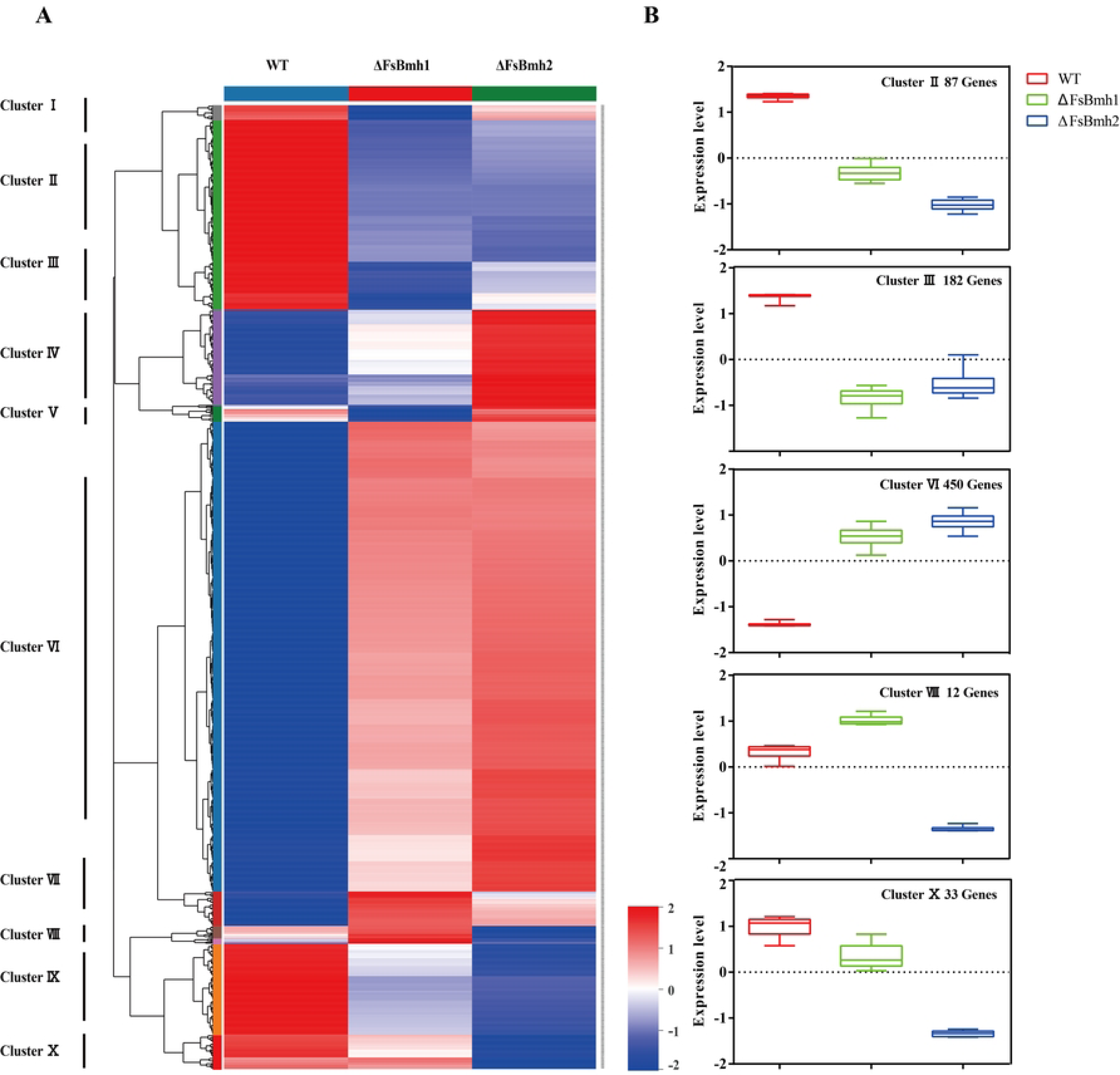
Hierarchical clustering showing the expression of WT, ΔFsBmh1 and ΔFsBmh2 DEGs. (A) A total of 981 DEGs were enriched in KEGG pathways (ΔFsBmh1 and ΔFsBmh2 versus WT), and gene expression pattern for 981 DEGs clustering into ten clusters. Red means that the gene is highly expressed, blue means that it is less expressed. (B) The distribution of gene expression values in clusters II, III, VI, VIII and X. Color scale showed the level of gene expression of *log*_10_^(FPKM+^ ^1)^.

**Fig 12.**
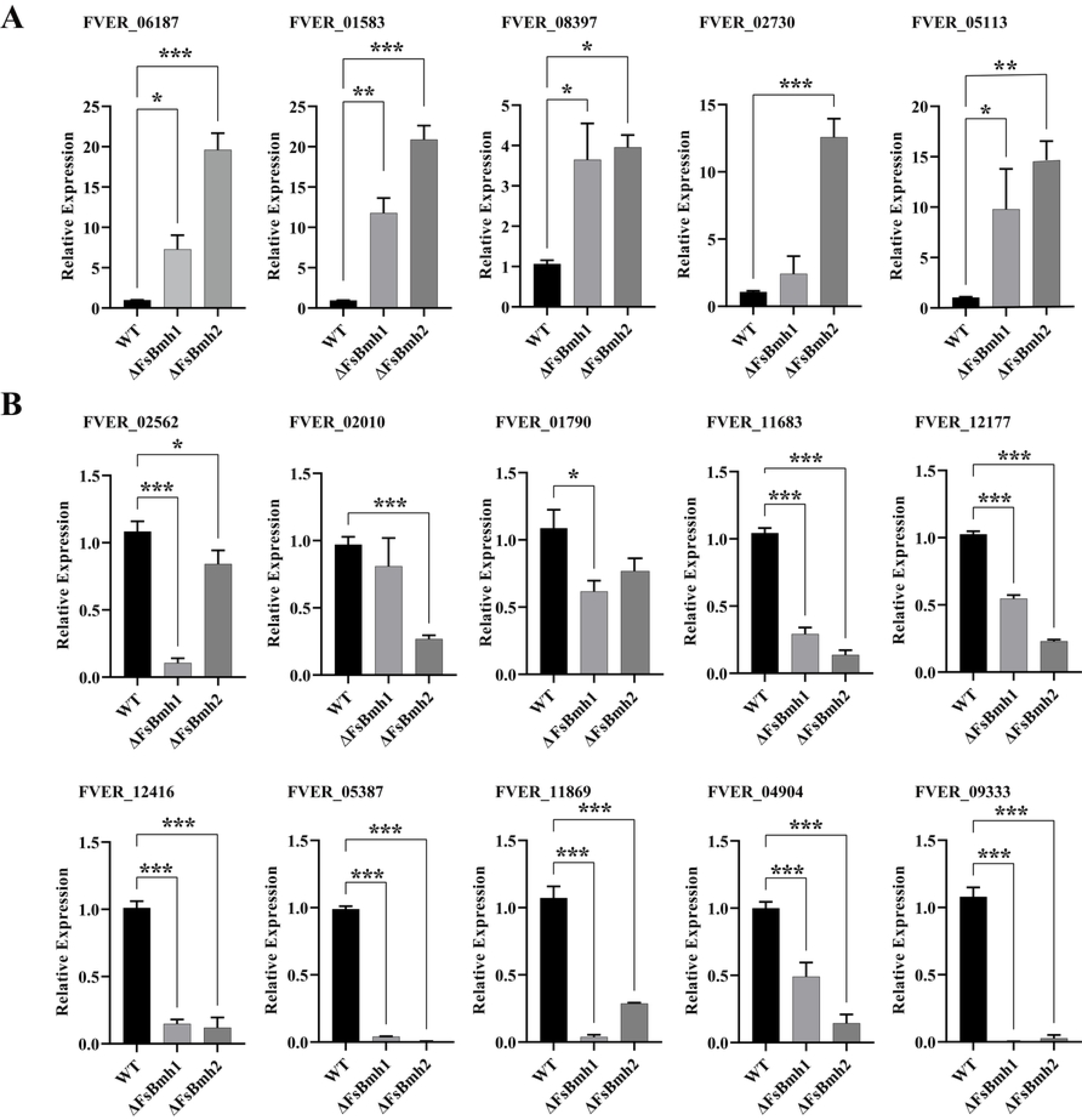
Validation of differentially expressed genes in ΔFsBmh1 and ΔsBmh2. (A) Up-regulated genes inΔFsBmh1 and ΔFsBmh2 versus WT. (B) Down-regulated genes inΔFsBmh1 and ΔFsBmh2 versus WT. These genes were associated with hyphal growth, sporulation and virulence The mean and standard deviation were derived from three independent biological replicates, each of which contained three technical repeats. The expression of each gene transcripts was measured by quantitative real-time RT-PCR(2*^−^*^ΔΔCT^ method). The relative transcript level was calibrated using 18S rRNA as reference, and the transcript levels of WT were set to a value of one. *P*-value were calculated based on Student’s T-test of replicate 2*^−^*^ΔΔCT^ values between a sample and WT. \**P<*0.05, \*\**P<*0.01, \*\*\**P<*0.001.

## Discussion

In this work, we investigated the role of *FsBmh1* and *FsBmh2* encoding 14-3-3 proteins in *F. sacchari*, a causal agent of sugarcane Pokkah boeng diease. Consistent with previous studies, 14-3-3 proteins are involved in multiple cellular processes in *F. sacchari*, such as hyphal growth, sporulation and virulence. Interestingly, we found that *FsBmh1* and *FsBmh2* regulate phenotypic traits in a reduntant manner but with distinct roles.

In previous reports, it was shown that *Bmh1* and *Bmh2* shared overlapping functions in regulate growth, conidiation, multistress responses, because deletion of either one of them exhibited similar defects in insect pathogenic fungus *Beauveria bassiana* and medicinal ascomycetous fungus *Ganoderma lucidm* [39, 40]. Compared to WT, ΔFsBmh-showed sparse aerial mycelium and lost the light brown pigment. Besides, the colony diameter and conidial yield of ΔFsBmh-were significantly decreased, but the defects of ΔFsBmh2 were more severe than ΔFsBmh1 (Fig 4A and Fig 4B). *AbaA*, *BrlA* and *WetA* are defined as the central regulatory pathway for conidiation [41]. Deletion *AbaA* or *WetA* specifically blocked conidia production and *BrlA* as a C2H2 zinc finger transcription factor acts upstream of *AbaA* and *WetA* [42, 43]. The transcriptional landscape of ΔFsBmh-showed that *AbaA* or *WetA* only strongly down-regulated in ΔFsBmh2 that is corresponding with previous results (Fig 2B and Fig 4B). Aside from their effects on growth and sporulation, *FsBmh1* and *FsBmh2* also play a pivotal role in pathogenicity. Either deletion *FsBmh1* or *FsBmh2* produced more slighter symptoms than WT (Fig 9). The phenomenon is different from *F. graminearum*, only *FgBmh2* contributed to virulence [44]. In order to futher clarify the functions of *FsBmh1* or *FsBmh2*, the overexpression mutants of *FsBmh*-were constructed. Intriguingly, a complementary relationship between *FsBmh1* and *FsBmh2* were observed. Deletion of one of the *FsBmh* genes resulted in upregulation of another *FsBmh* gene in the deletion mutants; similarly, overexpression one of the *FsBmh* gene resulted in downregulation of another *FsBmh* gene (Fig 3 and Fig 5A). These results suggest that there may be a sophisticated feedback mechanism to balance *FsBmh1* and *FsBmh2* in the cells. Furthermore, the deletion/silencing mutants ΔFsBmh1/sFsBmh2 and ΔFsBmh2/sFsBmh1 showed more severe manifestations to the fungus in general, suggesting an accumulation in function of the two 14-3-3 genes for virulence regulation (Fig 6). These phenomenon could be explained by the fact thatΔFsBmh1 andΔFsBmh2 shared up to 25.6% of DEGs in *F. sacchari* transcriptome that are associated with multiple phenotypes by regulating similar pathways (Fig 10).

Exceptionally, we noticed that *FsBmh1* and *FsBmh2* played distinct roles in regulating conidial germination and hyphal branching. Unlike ΔFsBmh1, deletion *FsBmh2* resulted in delayed germination. These results were conflict with *Beauveria bassiana* that conidial germination was equally accelerated in two 14-3-3 gene deletion mutants. The mechanism can be explained by *FsBmh2*-specific transcriptomes in cluster X (Fig 11B), the transcript level of *Chs1* and *BgIB* were decreased encoding chitin synthase 1 and -glucosidase respectively. They have been reported regulate conidial germination by cell wall remodeling in other filamentous fungi [45, 46]. Notably, only ΔFsBmh2 displayed more hyphal branches than the wild type strain and ΔFsBmh1/sFsBmh2 appeared more hyphal branches than ΔFsBmh1. These results indicated that *FsBmh2* negatively regulate hyphal branching. Moreover, we observed that *FsBmh1* and *FsBmh2* regulated conidia phenotypes in varied ways. On nutrient-rich medium, deletion of *FsBmh2* resulted in smaller microspores but deletion of *FsBmh1* resulted in thinner and longer microspores (Fig 6A and Table 2). However, on CLA medium use to induce the production of macrospores, we found that deletion of *FsBmh1* produced more macrospores and the ability of disruption of *FsBmh2* mutant strains were partially destroyed (Fig 7). Unlike other fungi, decreased transcript level of *FsBmh2* than *FsBmh1* resulted in shorter mycelial cell. This phenomenon is supported by the drastic upregulation of 3 genes (*Bud3*, *Bud4* and *SepH*) required for the control of hyphal septation in *FsBmh2*-specific transcriptomes [47]. Additionally, we noticed the expression of *Cdc25* and *Cdc45* in ΔFsBmh2 were in significantly higher level than in ΔFsBmh1, implying an improved rate of entering mitosis by regulating G1/S and G2/M transition, to promote mitosis [48–52]. It was recently reported that FgPMA1, a plasma membrane H+-ATPase specially interacts with FgBmh2 regulates pathogenicity in *F. graminearum* [53]. Whether similar mechanism present in *F. sacchari* remains to be investigated.

## Supporting information

**S1 Fig. Sequence alignments and analysis of the phylogenetic relationships among 14-3-3 and its homologs.** (A) sequence alignment of 14-3-3 and its homologs. (B) A neighbor-joining tree was constructed using the amino acid sequence of 14-3-3 proteins from different organisms.

**S2 Fig. Generation of *FsBmh1* deletion mutants and complementary strains.** (A), (B) *FsBmh1* gene locus and gene replacement construct. (C), (D) *FsBmh1* deletion mutants and complementary strains were validated by PCR with FsBmh1-A/B. (E) *FsBmh1* deletion mutants and complementary strains were validated by Southern Blot. The probes have been marked in the Figure.

**S3 Fig. Generation of *FsBmh2* deletion mutants and complementary strains** (A), (B) *FsBmh2* gene locus and gene replacement construct. (C), (D) *FsBmh2* deletion mutants and complementary strains were validated by PCR with FsBmh2-A/B. (E) *FsBmh2* deletion mutants and complementary strains were validated by Southern Blot. The probes have been marked in the Figure.

**S4 Fig. Generation of *FsBmh*-deletion/silenced mutants.** (A), (B) Silenced *FsBmh1* in ΔFsBmh2 and silenced *FsBmh2* in ΔFsBmh1. *FsBmh*-RNA interference vector was generated by insertion of the *FsBmh*-RNAi cassette using in-fusion strategy (see Materials and Methods). The mean and standard deviation were derived from three independent biological replicates, each of which contained three technical repeats. The expression of *FsBmh*-transcripts was measured by quantitative real-time RT-PCR(2*^−^*^ΔΔCT^ method). The relative transcript level was calibrated using 18S rRNA as reference, and the transcript levels of WT were set to a value of one. * indicates a statistically significant differences (*P<*0.05), ** indicates a statistically significant differences (*P<*0.01).

**S5 Fig. Generation of *FsBmh* - overexpression mutants.** (A), (B) overexpression *FsBmh1* and *FsBmh2* in wide type. *FsBmh*-overexpression vector was generated by insertion of the *FsBmh*-genes using in-fusion strategy (see Materials and Methods). The mean and standard deviation were derived from three independent biological replicates, each of which contained three technical repeats. The expression of *FsBmh*-transcripts was measured by quantitative real-time RT-PCR(2*^−^*^ΔΔCT^ method). The relative transcript level was calibrated using 18S rRNA as reference, and the transcript levels of WT were set to a value of one. * indicates a statistically significant differences (*P<*0.05), ** indicates a statistically significant differences (*P<*0.01).

**S6 Fig. FsBmh1 and *FsBmh2* influence hyphal growth by regulating cell length.** Hyphal of the wide type and all mutant strains were harvested after 8 h of incubation in PDB medium, at 28°C,150 rpm. Hyphal separation was stained with CFW and examined by fluorescence microscopy. Scale bar= 50 *µm*.

**S1 Table. Paired primers used for the manipulation of *Bmh1* and *Bmh2* in *F. sacchari* and the identification of their mutants.**

**S2 Table. Paired primers used in qRT-PCR for assessing transcript levels of 15 genes.**

**S3 Table. Grading standard of Pokkah boeng disease of sugarcane.**

**S4 Table. KEGG enrichment analysis of up-regulated DEGs between** Δ**FsBmh1 and WT.**

**S5 Table. KEGG enrichment analysis of down-regulated DEGs between** Δ**FsBmh1 and WT.**

**S6 Table. KEGG enrichment analysis of up-regulated DEGs between** Δ**FsBmh2 and WT.**

**S7 Table. KEGG enrichment analysis of down-regulated DEGs between** Δ**FsBmh2 and WT.**

**S8 Table. List and annotation of DEGs in five representation clusters.**

## Author Contributions

Conceived and designed the experiments: YZ, ZC, and CB. Performed the experiments: CY, ZL. Analyzed the data: CY, YZ, ZC. Wrote the paper: ZC, CY, YZ, and CB.

